# Mutational profiling of lipomas

**DOI:** 10.1101/585307

**Authors:** Deepika Kanojia, Pushkar Dakle, Anand Mayakonda, Rajeev Parameswaran, Mark E Puhaindran, Victor Lee Kwan Min, Vikas Madan, H Phillip Koeffler

## Abstract

Lipomas are benign fatty tumors with a high prevalence rate, mostly found in adults but have a good prognosis. Until now, reason for lipoma occurrence not been identified. We performed whole exome sequencing to define the mutational spectrum in ten lipoma patients along with their matching control samples. We identified 412 somatic variants including missense mutations, splice site variants, frameshift indels, and stop gain/lost. Kinase genes and transcriptions factors were among the validated mutated genes critical for cell proliferation and survival. Pathway analysis revealed enrichment of calcium, Wnt and phospholipase D signaling in patients. Whole exome sequencing in lipomas identified mutations in genes with a possible role in development and progression of lipomas.

**Author Summary:** We presented genomic insight into the development of lipomas, the most common benign tumor of soft tissue. Until date, no one knows the cause of lipoma development and its progression. Our group for the first time profiled ten lipoma patients’ samples and their matching normal controls to delineate the somatic mutation pattern using whole exome sequencing. Interestingly, exome-profiling results highlighted the potential role of important kinase genes and transcription factors in lipoma development. In addition, calcium, Wnt and phospholipase D signaling pathway speculated to be involve in pathogenesis of this disease.

## Introduction

Lipomas, benign soft tissue fatty tumors, are the most common benign mesenchymal tumors [1]. Lipomas are slow growing tumors and often occur under the skin on the neck, shoulders, arms, back, abdomen and thighs. However, occasionally lipomas may be present in deep location or originate within muscle [2]. Lipoma affects only 1% of the population, although it is probably underreported [3]. These benign tumors are commonly found in adults and their prevalence is higher in men than in women. Lipomas are generally small round masses of less than 5 cm and typically are harmless unless they compress an organ. Patients may have either a solitary lipoma or multiple lipomas. Lipomas commonly are diagnosed by clinicians and histologically examined after surgical resection. Interestingly, prognosis for lipomas is very good with almost no chance of recurrence after removal of the tumor [4].

Exact cause of lipomas is still not known, but lipomas have been identified with genetic rearrangements including structural changes at 12q13-15, 13q and 6p21-23 regions [5]. Approximately, 55%-75% of solitary lipomas have cytogenetic abnormalities involving *HMGA2* gene rearrangements (truncation or fusion) [6]. Studies revealed enhanced adipogenesis and a normal rate of adipocyte apoptosis in lipoma tissue compared to normal fat tissue [7].

A more aggressive form of fatty tumor is liposarcoma, which is rare in occurrence but they frequently develop dangerous metastatic tumors which is symptomatically different from lipomas [4]. Whole exome sequencing analysis on different subtypes of liposarcoma identified driver somatic mutations including PI3KCA, TP53, CHEK2, NF1, ATM, BRCA1 and EGFR [8]. Genomic characterization of tumors improved our understanding of their molecular genesis for better management of the disease.

We screened superficial subcutaneous lipomas using whole exome sequencing to systematically profile their mutational signatures, and identify affected pathways to understand further their molecular pathogenesis.

## Results and discussion

We performed a genomic analysis to profile the somatic mutational landscape of lipomas by whole exome sequencing. To best of our knowledge, this is first study to perform exome sequencing on lipomas.

### Clinical characteristics of lipoma patients

Primary subcutaneous lipomas were obtained from patients with their consent undergoing surgical resection at the National University Hospital. Detailed clinical features associated with these benign tumors presented in Supplementary Table 1. We collected lipomas from ten patients (all males) with a median age of 52 years. We performed whole exome sequencing on ten lipomas and normal controls from the same patient (either blood sample or adjacent normal tissue).

### Sequencing data

Whole exome sequencing of lipoma samples achieved an average of 135 million reads per sample (range 111-188 million). The control samples from the same patients found to have an average of 128 million reads per sample (range 112-164 million). Sequencing obtained a mean depth of 93 and 85 across the sequencing regions in lipomas and matching controls, respectively.

Variant allele frequency (VAF) implicate how early a mutation occurred during progression of the tumor and our data suggest that the mutations occurred both early (variants with high VAF) and later (variants with low VAF) during development of their lipoma. In total, 5 frameshift deletions and 5 frameshift insertions occurred with 5-20% VAF.

### Mutational landscape in lipomas

Using our data pipeline analysis, we identified 412 somatic variants with a median of 42 per patient sample (range 11-79) [Fig 1A and Supplementary Tables 2-11]. The type of mutations include 367 missense mutations, 19 stop gained, 5 stop lost, 9 splice acceptor, 2 splice site donor, 5 frameshift insertions and 5 frameshift deletions [Fig 1B]. Details of types of mutation for each sample listed in Supplementary Table 12. Similar to other benign tumors and sarcomas, overall mutation burden in lipomas observed to be relatively low. A total of 42 variants were confirmed to be mutated by Sanger sequencing including 37 missense mutations, 1 frameshift deletion, 2 in-frame deletions, 1 splice donor and 1 stop gained mutation. Detailed list of variants, their mutation type and known function of genes provided in Supplementary Tables 13. Validated mutations included kinase genes; ACVRIC (VAF 28%), PTPRT (VAF 14%), ERBB2 (VAF 14%) and MAPK7 (VAF 20%) and transcription factors ATMIN (VAF 7%), ZNF317 (VAF 26%) and FOSL1 (VAF 19%); each are crucial for cell survival and proliferation. No single mutated gene was common in all ten tumors. Several of the genes are known to play functional role in oncogenesis such as APC (VAF 26%), FOSL1 (VAF 19%), ERBB2 (VAF 14%), NFATC3 (VAF 10%), PDE1A (VAF 11%), ATMIN (VAF 7%), and MAPK7 (VAF 20%). In addition, experimental evidence have also indicated modulation of FOSL1 (VAF 19%) [9], ERBB2 (VAF 14%) [10], and PDE1A (VAF 11%) [11] regulates adipogenesis.

**Fig. 1.**
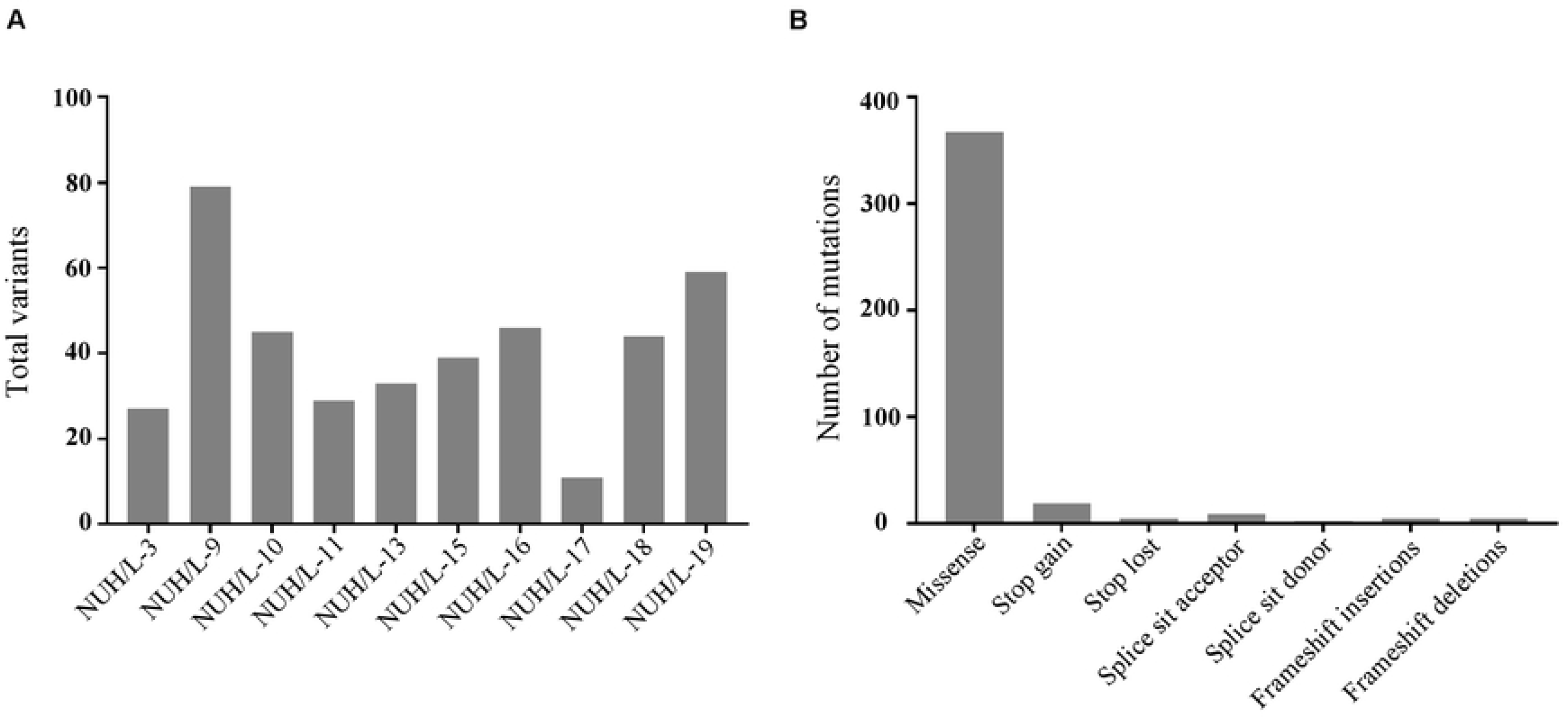
Variants identified in ten lipomas using whole exome sequencing. **(A)** Total number of variants found in each lipoma sample. **(B)** Number of each type of variant identified in ten lipoma samples.

### Mutational patterns in lipoma patient samples

We performed mutational spectrum analysis of lipomas to categorize their mutational signature and to identify functional mutagenic processes in lipomas. Analysis revealed mutational signature 15 as the major signature affecting lipoma patients [Figs 2A and 2B]. Signature 15 is one of the mutational signature associated with defective DNA mismatch repair. Patient wise signature analysis is shown in Supplementary Fig1.

**Fig. 2.**
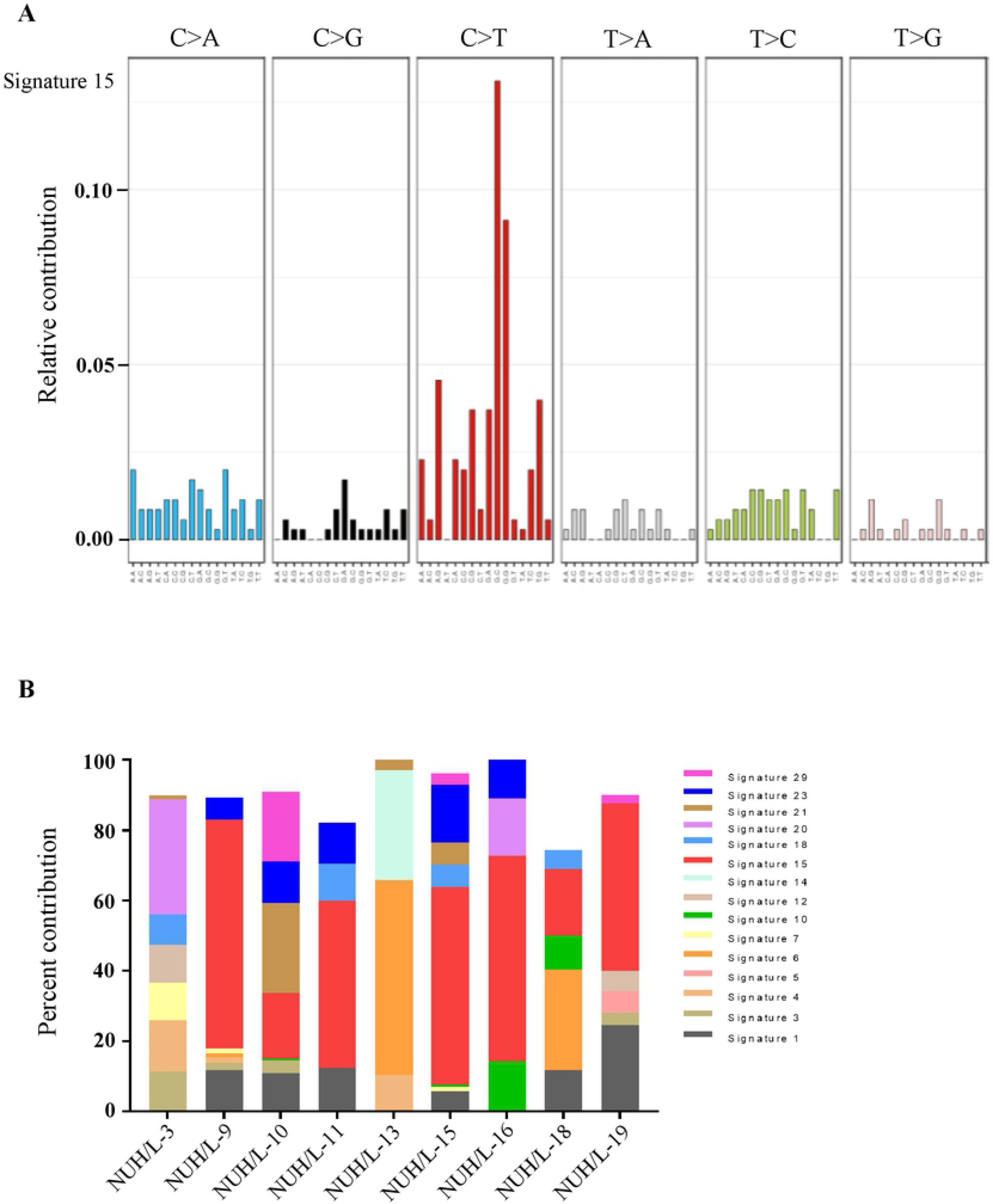
Signature of mutational process in lipomas. **(A)** Mutational signatures presented according to 96-substitution classification highlighting signature 15 as major process associated with lipomas. **(B)** Histogram shows percentage of signatures present in each lipoma sample.

### Altered pathways in lipomas

Pathway analysis was conducted to identify pathways and biological processes affected by mutated genes in lipomas. This analysis included gene variants validated and confirmed with Sanger sequencing. KEGG analysis indicated that out of 42 validated genes, eleven were involved in signalling of calcium (*P* = 0.0005), Wnt (*P* = 0.003), phospholipase D (*P* = 0.03) as well as stem cells’ pluripotency (*P* = 0.003) and axon guidance (*P* = 0.03) pathway [Supplementary Fig2A]. In addition, biological processes were identified which were associated with genes mutated in lipoma samples including calcium ion transport and microtubule regulation as well as signaling through the insulin receptor and tyrosine kinase [Supplementary Fig2B]. Interestingly, calcium signalling pathway and calcium ion transport biological process emerged as a major pathway/process affected by mutated genes in lipoma. Several studies have indicated a regulatory role of calcium ions in adipogenesis contributing to increased fat tissue by promoting and accelerating pre-adipocyte differentiation [12, 13], which is a feature of lipomas. In addition, Wnt signaling [14] and phospholipase D signalling [15] pathways shown to regulate adipogenesis.

## Conclusions

We identified and validated mutations in ten lipoma patient samples using whole exome sequencing analysis. However, recurrent gene variants were not noted in ten lipoma patients; thus these alterations could be passenger mutations. Additional samples need to be sequenced and the functional consequences of the identified mutations need to be explored to support the present findings. Present analysis also delineated lipoma-specific mutational signatures and identified wide-ranging genomic landscape of altered genes and pathways in lipomas that could be useful to investigate further the functional cause of lipoma and its progression.

## Materials and Methods

### Tissue collection and sample preparation

Lipomas, normal adjacent control tissues and patient’s blood samples were obtained from National University Hospital, Tissue repository, Singapore in accordance with ethical guidelines of NUH Institutional Review Board. According to their requirement, samples were collected after obtaining informed written consent from patients. Tissue samples or blood received after surgery was snap frozen and stored in liquid nitrogen until processing. Genomic DNAs were isolated from tumors, control tissues and blood samples using QIAamp DNA Mini Kit (Qiagen) according to the manufacturer’s instructions. Genomic DNAs were quantified by Qubit dsDNA BR assay kit (Life technologies) and their quality verified by agarose gel electrophoresis.

### Whole exome sequencing

Exome libraries were prepared using the Agilent SureSelect All Human Exon v5 capture kit [8]. Briefly, the qualified genomic DNA samples were randomly fragmented into base pair peak of 150 to 200bp, and adapters ligated to both ends of the resulting fragments. Adapter-ligated templates were purified by the Agencourt AMPure SPRI beads, and fragments with insert size of about 200 bp were excised. Extracted DNA was amplified by ligation-mediated polymerase chain reaction (LM-PCR), purified, and hybridized to the SureSelect Biotinylated RNA Library (BAITS) for enrichment. Hybridized fragments were bound to the streptavidin beads whereas non-hybridized fragments were washed away after 24h. Captured LM-PCR products were subjected to Agilent 2100 Bioanalyzer to estimate the magnitude of enrichment. Each captured library was independently loaded on Hiseq4000 platform for high-throughput sequencing to ensure that each sample meets the desired average fold-coverage. Raw image files were processed by Illumina base calling Software 1.7 with default parameters, and the sequences were generated as 90bp paired-end reads.

### Analysis of whole exome sequencing

Paired end sequencing reads were aligned to the human reference genome (hg19) using bwa mem. Samblaster was subsequently used to mark PCR duplicates [16]. Systematic errors in base quality scores were then corrected using GATK4 Base Recalibrator [17]. Somatic single nucleotide variants and short indels were detected against a matched normal using a consensus calling approach with both GATK4 Mutect2 and Strelka2 [18]. Variants calls from Mutect2 further filtered with GATK4 FilterMutectCalls. Vcf2maf was used to convert the. vcf files to MAF format and also annotated using Ensembl VEP (Version 92_GRCh37) [19]. Depth of coverage for all samples calculated using GATK3 DepthOfCoverage.

We excluded any variants which were not tagged PASS by FilterMutectCalls3 and Strelka2. Variants were further filtered to remove those with global minor allele frequency (GMAF) lower than 1% or variants tagged as common variants by vcf2maf based on allele count in Exac populations, while still retaining those classified as “pathogenic” in ClinVar. Variants with either LOW/MODIFIER impact as per VEP or with variant allele frequency (VAF) less than 0.01 were also removed.

Each mutation categorized into seven categories AT transitions, AT transversions, CpG transitions, CpG transversions, CG transitions/ transversions and Indels. MuSiCa, online tool was used for somatic characterization of mutation signatures present in the patient’s variants list [20]. Refseq-annotated mutations (genes) were used for pathway analysis using PathScan algorithm integrated into MuSic pipeline [21, 22].

### Sanger sequencing validations

Gene variants with VAF greater than 10% were validated by Sanger sequencing. Polymerase chain reaction was performed using 25 ng template DNA of lipoma tissue and normal control sample from the same individual. PCR products run on 1% agarose gel and purified using Wizard^®^ SV Gel and PCR Clean-Up System (Promega). Purified PCR products were sequenced and analyzed with the Sequence Scanner Software 2 Version 2 (Applied Biosystems).

## Declarations

### Availability of data and materials

All data generated during this study are included in the published article and its supplementary tables.

### Authors’ contributions

DK and HPK designed the experiments; DK performed the experiments; DK, DPP, AMT and VM analyzed the results; RP, MEP and VLK provided the patient samples; HPK supervised the project and DK, DDP and HPK wrote the manuscript. All authors read and approved the final manuscript.

### Ethics approval and consent to participate

Patient’s tissues and blood samples collected in accordance with ethical guidelines of NUH Institutional Review Board and after obtaining informed written consent from patients.

### Consent for publication

Not applicable.

---

**Supplementary Fig. 1:** Mutation signature associated with each lipoma patient.

**Supplementary Fig. 2:** Enriched pathways (A) and biological processes (B) identified and associated with validated mutated gene products in lipoma tumors.

